# Structural influence of the conserved Hsp40 HPD tripeptide on Hsp70 chaperone function

**DOI:** 10.1101/2020.03.25.008359

**Authors:** OO Mallapre, NAD Bascos

## Abstract

The control of Hsp70 functions has been related to the modulation of ATP hydrolysis and substrate capture by Hsp40. Structural and biophysical analyses of Hsp40 variants and their interactions with Hsp70 have identified key residues for this functional control mechanism. Conserved residues in both Hsp40 and Hsp70 have revealed conserved interactions that link Hsp40 binding to the catalytic residues within Hsp70. The current work investigates the effect of documented J-domain dysfunctional mutations (i.e. D35N, H33Q) on the described interaction linkage. Molecular dynamics simulations were used to compare the persistence of individual bond types (i.e. H-bonds, salt bridges, hydrophobic interactions) between Hsp70 and the bound forms of functional and dysfunctional Hsp40 variants. The generated data suggests the involvement of both direct and allosteric effects for the tested mutations. The observed changes relate mutations in the conserved HPD tripeptide of Hsp40 to alterations in the interaction network that induces Hsp70 chaperone functions.

**STATEMENT OF SIGNIFICANCE:** The significance of the work may be summarized as follows. First, the interaction network for the simulated systems were observed to be different from one previously proposed for a disulfide linked complex (9). This may be attributed to altered residue movement and interactions without the restrictions set by the disulfide link. These results support the use of *in silico* methods to refine investigations of molecular contacts, particularly for systems, whose in *vitro* structural elucidation are difficult to achieve without modifications.

Second, key interactions for intermolecular and intramolecular contacts were observed within a short simulation time (0.1 ns) matched those from much longer runs (500 ns) (4). This result highlights the possibility of identifying key interactions with relatively low computational cost.

## INTRODUCTION

The Hsp70 family of proteins consists of molecular chaperones that are extremely well-conserved among all domains of life. Despite being named heat shock proteins, Hsp70s also normally function under non-stressful cell conditions. As molecular chaperones, Hsp70s have been implicated in the proper folding of nascent polypeptide chains, the prevention of protein aggregation, the transport of polypeptides across membranes, the refolding or remodeling of mature polypeptides and the facilitation of protein degradation (1). These functions are made possible through a repeating cycle of substrate-binding and release regulated by ATP hydrolysis and nucleotide exchange. In the ATP-bound state, the chaperone adopts an open configuration for its substrate-binding domain (SBD), exposing a hydrophobic patch for substrate binding, while assuming a docked conformation with the nucleotide-binding domain (NBD). After ATP hydrolysis, the molecule shifts into a closed SBD configuration where the substrate binding domain beta sheet region (SBDβ) clamps around the substrate and undocks from the NBD. The SBD and NBD remain attached through a short linker that has been implicated in modulating the domain interactions. In the ATP bound state, the linker has been observed to bind the NBD facilitating the docked conformation (2, 3). Disruption of the NBD-linker interaction has been suggested to promote the undocked conformation associated with ATP-hydrolysis (4). Exchanging the bound ADP with ATP returns the chaperone into the open configuration which restarts the cycle (5).

Hsp70’s activity cycle is facilitated by its cochaperone, Hsp40, which promotes ATP hydrolysis and substrate capture, and a nucleotide exchange factor, which replaces bound ADP with ATP. The Hsp40 family is composed of a structurally diverse set of proteins found in various organisms like *E. coli, S. cerevisiae* and humans. Despite their structural diversity, they all share a well-conserved J domain (Jd), which has been found to be crucial for their interaction with Hsp70s (5). Altered interactions between the Hsp40 Jd and Hsp70s, both in terms of binding affinity (6, 7), and mechanical properties (8), have been related to changes in the ability of Jd to promote Hsp70 functions. A recent report on the structure of a Jd docked to an Hsp70 molecule in *E. coli* revealed a network of polar and nonpolar interactions proposed to transmit the signal from the Jd to the catalytic center at the NBD (9). This network involves residues (in both Jd and Hsp70) that are important for Hsp70 function and whose mutation are proposed to abolish activity for the Hsp40-Hsp70 molecular machine. Two mutations, D35N and H33Q are located at the conserved HPD tripeptide of Jd (6), and are well documented to disrupt Hsp70 function (6,7). This study aims to investigate the nature of the disruption caused by these two mutations in the context of the proposed Hsp40-Hsp70 interaction network in *E. coli* (9).

## MATERIALS AND METHODS

A structure of full-length *E. coli* DnaK bound to the J domain of DnaJ was obtained from the Research Collaboratory for Structural Bioinformatics Protein Data Bank (RCSB PDB ID: 5NRO). The structure contained sequence modifications that aided crystallization (i.e. a T199A mutation to inactivate the enzyme (11), E47C and F529C mutations to stabilize the open conformation with a disulfide bridge, and replacement of the unstructured C-terminal with 5xHis). These were reverted to restore the protein’s native sequence (i.e. primary structure). The missing C-terminal residues were not rebuilt. The model with this restored sequence was used as the wildtype sample. The two mutants were generated by replacing the appropriate amino acid with the mutated version. Each sample, consisting of structures of the proteins (DnaJ and DnaK), ATP, and the cofactor Mg^2+^, was immersed in a rectangular water box using the Solvate package of Visual Molecular Dynamics (VMD). This placed a 5 Å padding of TIP3p water molecules around the protein structure.

Molecular dynamics simulations were performed with Nanoscale Molecular Dynamics (NAMD) 2.10 using the Combined CHARMM All-Hydrogen Parameter File (CHARMM22 All-Hydrogen Parameter File for Proteins and CHARMM27 All-Hydrogen Nucleic Acid Parameter File) force field. The Particle Mesh Ewald Sum method was used to deal with electrostatic interactions. The water-immersed structures were minimized for 100 steps of the default conjugate gradient and line search algorithm then brought to a temperature of 310 K by assigning random velocities to all atoms according to a Maxwell distribution. Simulation ran in NPT environment with a time step of 2 fs for 50,000 steps, simulating 100 ps of molecular movement. Each sample was simulated thrice starting from the same water-immersed structure (12).

Predicted changes in the protein structures were analyzed through several parameters. Shifts in mobility were estimated based on Root Mean Square Deviation (RMSD) using the first frame as reference, where samples have an average RMSD of 0.584 Å. RMSD of the alpha carbon was calculated for each residue and for each frame of the simulation using a custom script. Energy from Van der Waals and electrostatic interactions between DnaK and DnaJ through time was measured using the NAMDEnergy plugin of VMD. H-bond occupancy was calculated for each possible residue pair using the Hydrogen Bonds plugin of VMD with a maximum donor-acceptor distance of 3.5 Å and an angle cut-off of 20°. Similarly, all possible salt bridges were identified using the Salt Bridges plugin of VMD with an oxygen-nitrogen distance cut-off of 3.2 Å. Occupancy was calculated with a cut-off distance of 9.0 Å (13). Finally, the occupancy of nonpolar interactions was calculated for each nearby pair of nonpolar sidechains using a custom script, with a distance cut-off of 4.0 Å, similar to that used in a previous study (9). Observations of differences in interaction between the wildtype Jd-bound, and the mutant Jd-bound DnaK systems were restricted to those with occupancy persistence differences of 30% or greater from the wildtype.

Using the list of residue pairs interacting through H-bonds, electrostatic and nonpolar interactions, the shortest path from the conserved HPD motif of Jd to the catalytic center, particularly residue K70, was traced for the wildtype and mutant versions of Jd, using a custom script. Only interactions with occupancies greater than 30% (i.e. bonds were persistent at least 30% of the simulation time points) were considered. Statistical analyses were conducted using Analysis of Variance (ANOVA) and Student’s T test, both with an α = 0.05.

## RESULTS AND DISCUSSION

### Residue RMSD

Molecular dynamics simulations were performed for models of the Hsp40-Hsp70 complex of *E. coli* (DnaJ and DnaK, respectively), with either wildtype or mutant (e.g. JdH33Q, JdD35N) versions of J-domain attached. The models were based on a deposited structure for the Hsp40-Hsp70 complex with a bound wildtype J-domain (PDBID: 5NRO) (9). Models for the mutant J-domain bound structures were made by the modification, and equilibration of the wildtype J-domain bound DnaK structure. The average root-mean-square deviation (RMSD) of each residue from each model was calculated for the whole duration of the simulation.

Comparing the residue RMSD of the mutants with the wildtype, most of the changes in RMSD, for both increased and decreased values, occur at the region close to active site of DnaK (Figure 1a & b). This suggests that the mutations in the J domain caused allosteric changes in mobility for residues around the active site of DnaK, possibly affecting the stability of interactions within the region.

**Figure 1.**
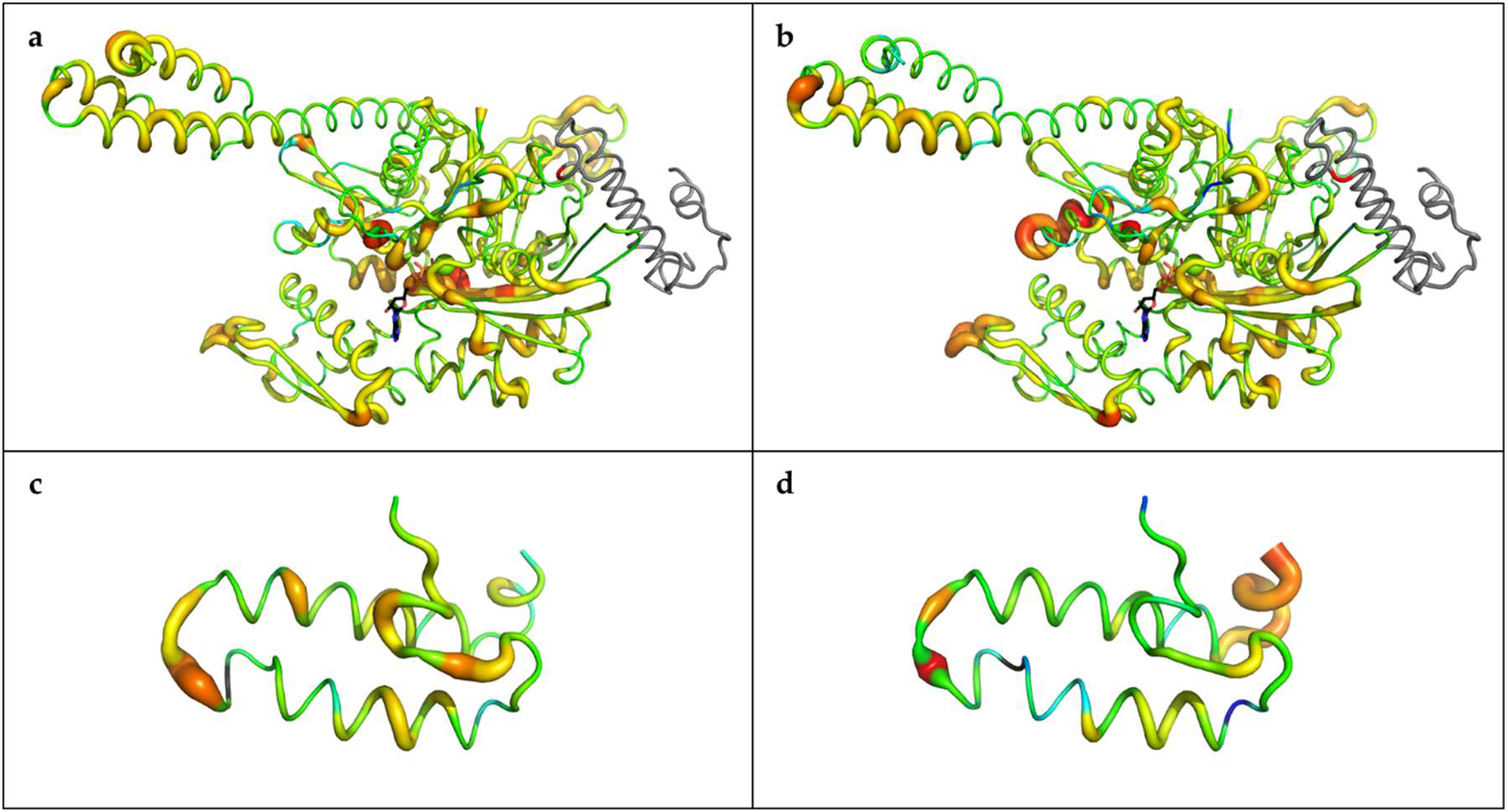
Mobility of residues in mutants relative to wildtype. The figure shows a tube representation of the mutants: (**a** and **c**) D35N; (**b** and **d**) H33Q. The thickness of the tube corresponds to the average mobility of the residue in the sample relative to the wildtype. Thicker tubes represent regions more flexible in mutants and thinner tubes represent regions less flexible in mutants. The tubes are colored by relative flexibility, increasing from blue to red, with green indicating residues of similar mobility between mutant and wildtype. The top panel (**a** and **b**) shows DnaK with Jd colored in gray, and the mutated residue colored red. The bottom panel (**c** and **d**) shows Jd alone with the mutated residue colored black.

The residues of Jd had very low RMSD compared to DnaK and relative changes in RMSD were not obvious in the putty representation when considering both chaperones. However, when the RMSD range is limited to those observed with Jd, changes become more apparent. In both mutants, an increase in mobility is observed at the terminal ends of Jd and at the loop between helix II and III. More pronounced changes were observed in D35N, where increases in mobility occur within helix II and III (Figure 1c & d). Helix II of Jd has been previously observed to bend upon binding DnaK, with the D35N mutant exhibiting increased bending (7). Although bending of helix II was not observed during the simulation, the observed changes in mobility of helix II in D35N could contribute to this phenomenon. This may also explain the lack of rigidity previously observed in the D35N mutant (8).

### Changes in interactions between residues

The changes in residue RMSD could be a result of rearrangements of bond networks within the two chaperones. To test this hypothesis, the H-bonds, as well as the electrostatic and nonpolar interactions were quantified based on bond occupancy. This is defined as the frequency of interaction between two residues throughout the simulation. Values are presented in percentage of simulation time for which the bond was observed. The nature and location of changes to this property may provide a rationale for the observed dysfunction of the Hsp40-Hsp70 system in the presence of the D35N and H33Q mutations in Jd.

The occupancies of some interactions were observed to vary significantly among the wildtype-Jd bound and mutant-Jd bound complexes (ANOVA, α = 0.05, n = 3). In those intermolecular interactions whose average occupancy changed by at least 30% from the wildtype complex, there was a significant difference across the three sets (Jd-DnaK; JdD35N-DnaK; and JdH33Q-DnaK). Given the same selection criteria, no significant difference was observed for the populations of intramolecular residue interaction occupancies within DnaK across the three sets.

This is evident in the contact map of DnaK shown in Figure 2. Despite the general similarity of the intramolecular contacts within the 3 Jd-bound systems, a limited number of big shifts in contact were observed suggesting that the effect of the selected Jd mutants on DnaK might occur through specific key locations, and small changes in residue configurations and interactions, which occur almost uniformly throughout the DnaK molecule.

**Figure 2.**
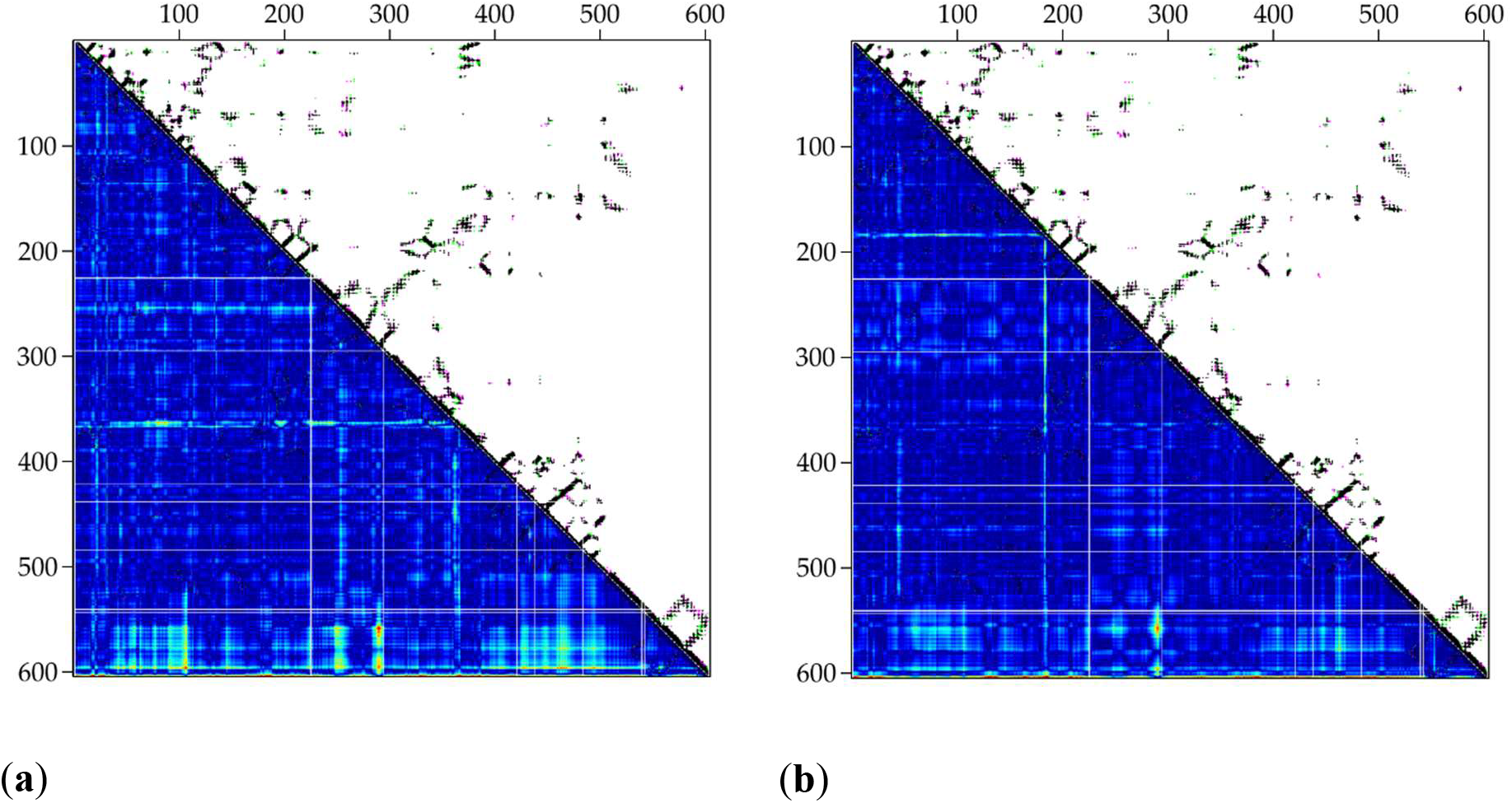
Map of all inter-residue contacts in DnaK. The x and y-axes both represent residue numbers while each point on the map represents a contact between two residues. The maps shown are superimpositions of the final timepoints of one replicate of the wildtype and each mutant: (**a**) D35N; (**b**) H33Q. On the upper-right side, black points represent contacts present in both wildtype and mutant, pink points represent contacts unique to the wildtype, and green points represent contacts unique to the mutant. The lower-left side is colored based on distances between alpha carbons of each residue in mutants, where blue regions are above the 8-Å cutoff, green regions are around the cutoff, and red regions are below the cutoff.

### Changes in Jd-DnaK Intermolecular contacts

#### Summary of altered contacts: D35N and H33Q

For this section and the rest of the discussion, values stated for occupancy are the average occupancy across three replicates. Pairwise comparison of each residue involved in intermolecular interactions revealed significant changes in occupancy for three H-bonds in the JdD35N-bound complex. These were interactions of Jd D/N35 with DnaK R167 (80.50% to 49.17%) and DnaK Q378 (64.5% to 14.17%), and Jd M30 with DnaK T215, which only decreased slightly (0.83% to 0%).

In H33Q, two H-bonds and one salt bridge significantly changed in occupancy. One of these H-bonds was D35-R167, but in contrast to the decreased occupancy with JdD35N, the JdH33Q complex showed an increase in occupancy from 80.50% to 92.17%. The second H-bond was with Jd Y25 and DnaK E217, where occupancy also increased from 3.00% to 55.50%. Meanwhile, the salt bridge between Jd R27 and DnaK D208 was reduced from 13.00% to 0.33%.

### Alterations with JdD35N

#### JdD/N35 with DnaK R167 (80.50% to 49.17%)

The D35-R167 bond has been previously reported to be a necessary interaction between DnaK and Jd (6). Recent work has suggested the need for conserving D35-R167 contact in Hsp40-Hsp70 systems across different species (4). This need may stem from the requirement to link J-domain association with the induction of ATPase activity in DnaK. The significance of changes in the D35-R167 interaction may lie in the allosteric changes observed in DnaK intramolecular contacts as discussed in the succeeding section.

#### JdD/N35 with DnaK Q378 (64.5% to 14.17%)

The reduction in occupancy between D/N35 and Q378 coincides with a significant decrease in occupancy between R167 and Q378, from 27.33% to 0.50%. Through its interaction with R167, Q378 is also known to stabilize the network of interactions in the DnaK catalytic site. Therefore, in addition to its interaction with R167, D35 could also attract Q378 and further stabilize the Q378-R167 interaction, enhancing signal transmission between the substrate and ATP. Loss of this key residue with the D35N mutation therefore results in the disruption of crucial interactions that allow Jd to stimulate ATP hydrolysis in DnaK.

#### Jd M30 with DnaK T215 (0.83% to 0%)

Residues T215 and E217 of DnaK’s NBD lobe II were previously reported to either bind DnaJ or stimulate hydrolysis. Their interactors, M30 and Y25 are both located on (Jd) helix II. A very small but significant reduction in M30-T215 was observed in D35N, while the opposite is observed in Y25-E217 in H33Q.

### Alterations with JdH33Q

#### Jd Y25 and DnaK E217(3.00% to 55.50%)

A significant increase in M30-T215 binding is observed in Y25-E217 in H33Q (3.00% to 55.50%). It must be noted that a small but significant increase in occupancy (from 1.67% to 4.33%) between E217 and L392 of the linker was also observed. This suggests that the effect of the change in Y27-E217 occupancy in H33Q mutants could propagate through the linker.

Increased affinity of the linker to the NBD could inhibit the conformational change to an undocked state associated with ATP-hydrolysis and substrate capture.

### Paired interactions: Jd R27-DnaK D208 (13.00% to 0.33%) and Jd D35-DnaK R167(80.50% to 92.17%)

The interaction between DnaJ R27 and DnaK D208 is the only intermolecular salt bridge observed to change occupancy in mutants, specifically in H33Q (from 13.00% to 0.33%). Analogues of these residues are observed in the Hsc20-Ssq1 complex of *S. cerevisiae* to be one of the predominant contributors to the interactions between the two molecules. This interaction, together with other electrostatic interactions were proposed to be involved in DnaJ recognition by DnaK. Mutation of even one of these residues was enough to reduce ATP hydrolysis in *S. cerevisiae* (4). Thus, the reduction of occupancy of R27-D208 in H33Q may contribute to a reduction of ATPase activity, perhaps through impairment of DnaJ recognition.

Our results in H33Q show a decrease in occupancy of R27-D208, paired with the increased propensity for D35-R167 (80.50% to 92.17%). These may represent a shift in the binding character of the J-domain with the H33Q mutation. Previous work has suggested the possibility for a range of interaction modes to be employed in Hsp40-Hsp70 systems across different species, provided that the key residue contacts are maintained (4). Disruption of these key interactions, as observed with altered bond propensities may contribute to the inability of the mutants to induce ATPase activity and DnaK function.

### Allosterically induced changes in DnaK intramolecular contacts

Interactions with both mutant J-domains generated allosteric changes in intramolecular contacts for DnaK residues. The most noticeable shifts in contact occur close to the C-terminal end (~ AA 550 – 604) of DnaK. These are seen with the increased green to red coloration of the alpha carbon distances. Notable changes were however observed in residue contacts associated with the DnaK NBD lobes, and between the NBD lobes and SBDβ. The intramolecular contacts altered by mutant Jd association varied depending on the partner protein (Figure 4).

**Figure 3.**
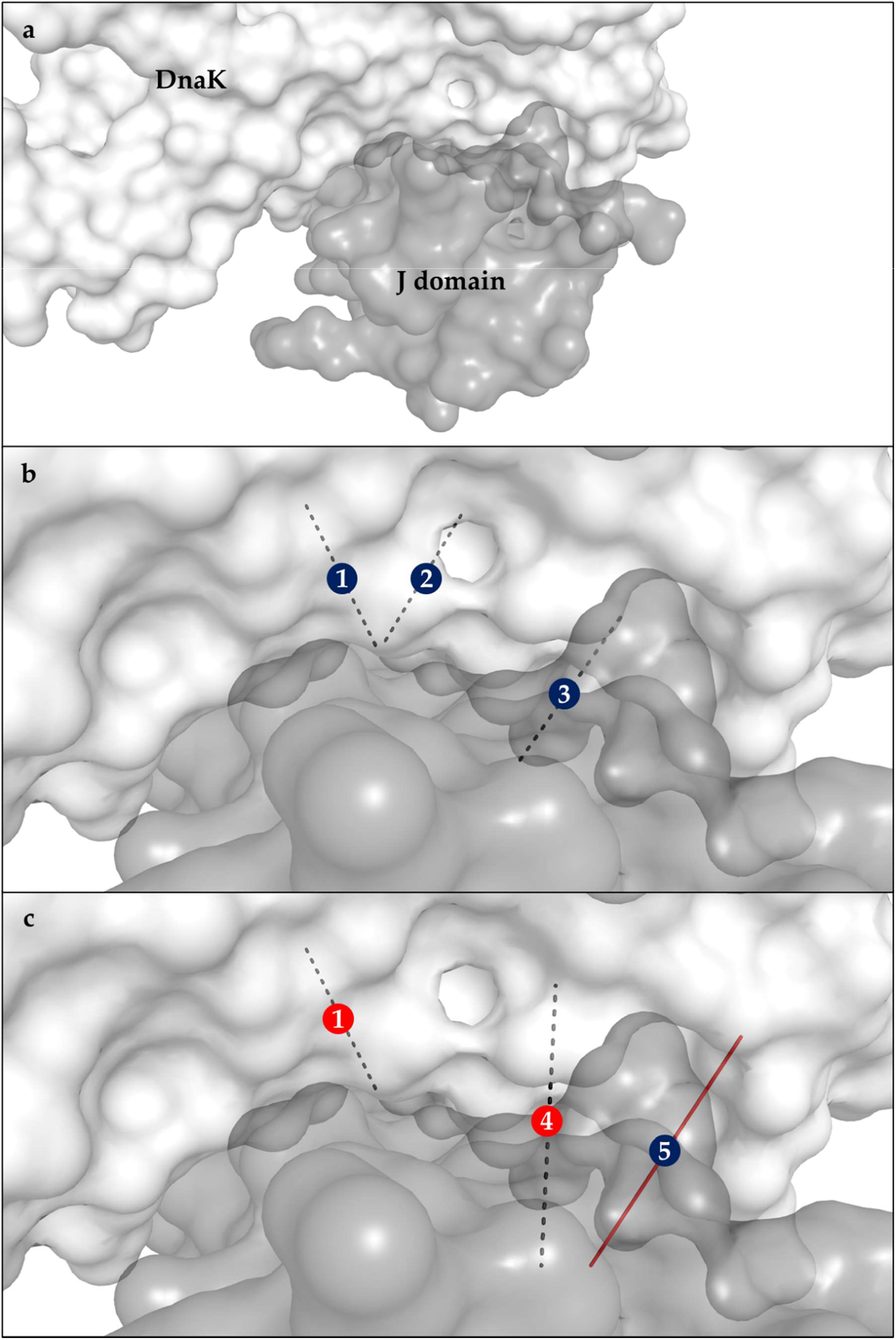
Interactions between DnaK and the J domain. The interactors are represented by translucent surfaces (DnaK: white, Jd: dark gray) while the interactions between them that significantly changed occupancy are shown as lines connecting the α-carbons of residues and colored based on type (H-bond: dashed, electrostatic: solid red, nonpolar: solid green): (a) The position of Jd relative to DnaK; A close-up view of interactions in: (b) D35N; (c) and H33Q. Labels: (1) R167-D35, (2) Q378-D35, (3) T215-M30, (4) E217-Y25, (5) D208-R27. Labels are colored by change in occupancy (increase: red, decrease: blue).

**Figure 4.**
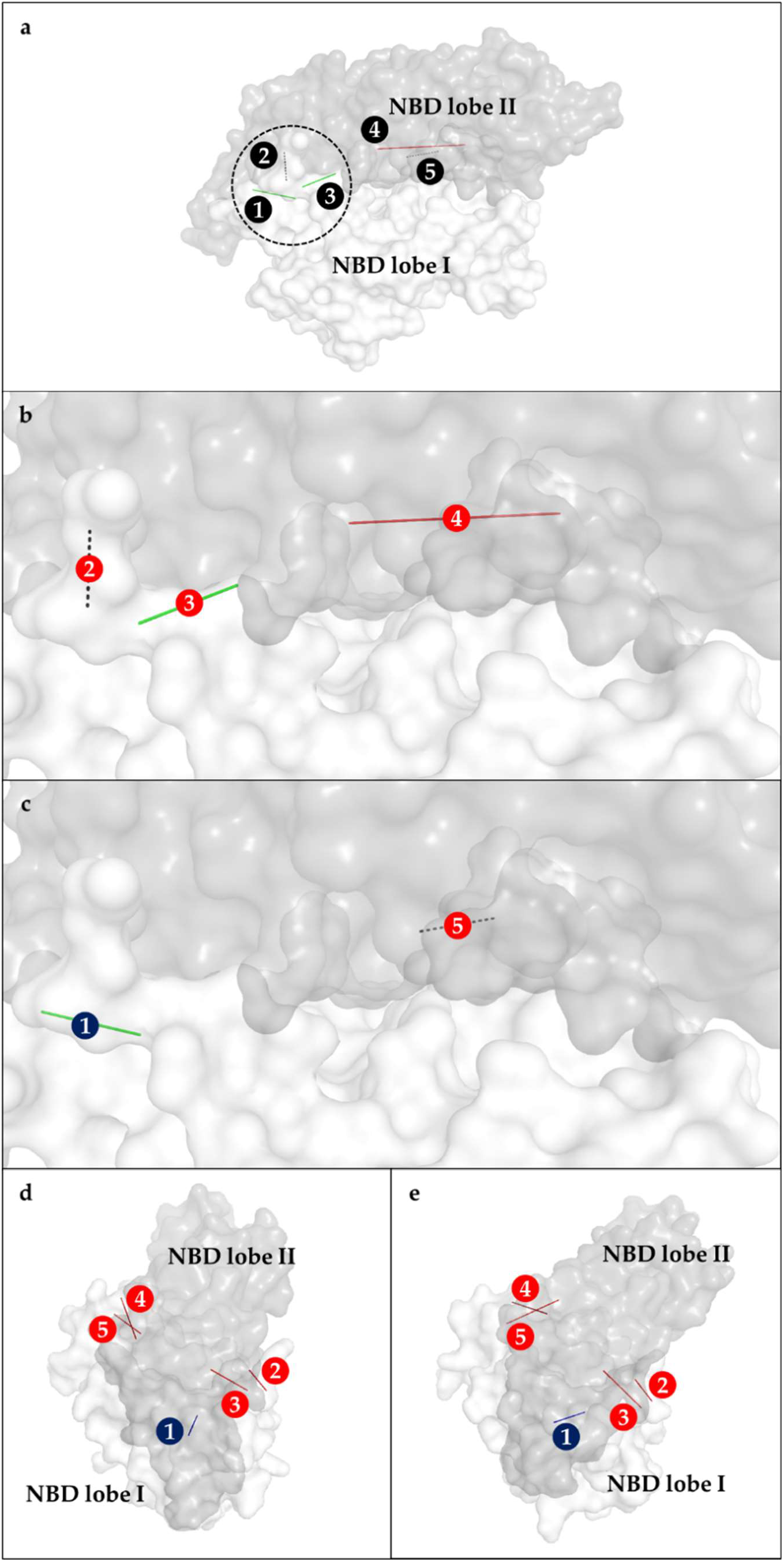
Interactions between the lobes of the nucleotide-binding domain. The interactors are represented by translucent surfaces (lobe I: white, lobe II: dark gray) while the interactions between them that significantly changed occupancy are shown as lines connecting the α-carbons of residues and colored based on type (H-bond: dashed, electrostatic: solid red, nonpolar: solid green): (**a**) The position of lobes relative to each other with the hinge region encircled; A close-up view of interactions that significantly changed occupancy in: (**b**) D35N; (**c**) and H33Q; A side view of the NBD with interactions colored by change in occupancy (increase: red, decrease: blue): (**d**) in ATP-bound open configuration; (**e**) and ADP-bound closed configuration. Labels: (1) A176-I207, (2) R362-D364, (3) V340-V365, (4) R84-D224, (5) E81-H226. Labels are colored by change in occupancy (increase: red, decrease: blue).

### JdD35N-DnaK specific allosteric changes

With JdD35N binding, several intramolecular bonds in DnaK were affected. The R362-D364 H-bond significantly increased in occupancy from 75.17% to 90.17% and the V340-V365 nonpolar interactions also significantly increased in occupancy from 31.33% to 66.17%. Both of these bonds are located in the hinge region of DnaK NBD lobe II. Interestingly, a significant increase in occupancy was also observed between residues in NBD lobe I (R84) and NBD lobe II (D224). The R84-D224 salt bridge connects both DnaK NBD lobes I and II and its occupancy varied from 8.17% to 59.83% between the wildtype Jd-bound, and the JdD35N-bound DnaK complexes.

An interaction between R167 in the NBD and D481 of the SBDβ is observed in the crystal structure of Jd-bound DnaK from *E. coli* (PDB ID: 5NRO) (9). This bond is proposed to be an important residue for communication between the two functional domains of DnaK. It is believed to inhibit ATPase activity in the NBD until triggered by substrate binding in the SBDβ (10).

The R167-D481 bond was preserved with JdD35N binding to have the same occupancy of 100% as the wildtype. In addition, another interaction with the SBDβ is observed with R167 and D479, which lies on the same loop as D481. This interaction had increased in occupancy in with JdD35N (43.83% to 100%). The additional contact with the SBDβ would help prevent the undocking of the substrate binding domain, suggesting an explanation for the inability of JdD35N to stimulate ATPase activity of DnaK.

Another interaction between NBD and SBDβ that changed in D35N was the D326-N415 H-bond. Occupancy significantly decreased from 38.33% to 4.83% with JdD35N binding. D326, together with its pair K414 is known to be important in interdomain communication between substrate and ATP, similar to D481. While the occupancy of the D326-K414 salt bridge did not change with JdD35N binding, the reduction in occupancy of the neighboring D326-N415 H-bond could indicate a weakening of interactions between the NBD and SBDβ at this location. This is interesting since the opposite effect is seen in a different side of the NBD where I167 connects with D479 of the SBDβ. In a proposed model of DnaK mechanism, I168-D481 and D326-K414 prevent the back-rotation of NBD lobes thus preventing premature ATP hydrolysis. Substrate association disrupts the I168-D481 interaction, allowing back-rotation of lobe I and ATP hydrolysis (10). An increase in interaction proximal to the I168-D481 bond (i.e. R167-D479) may therefore inhibit the movement of lobe I and ATPase function.

The induction of greater bonding propensity for I168-D481, but not D326-K414 by the JdD35N mutant results in an imbalanced shift that does not achieve the closed conformation associated with ATP hydrolytic activity.

### JdH33Q-DnaK specific allosteric changes

JdH33Q binding induced several intramolecular bond changes in the DnaK. Increased occupancy is observed for the R167-D479 interaction (43.83% to 93.17%) which connects NBD lobe I and the SBDβ; and the E81-H226 H-bond (0.50% to 3.33%) between the NBD lobes I and II. Decreased occupancy is observed for the A176-I207 non-polar interaction near the hinge of the two NBD lobes (81.5% to 47.3%). The increased occupancy seen with R167-D479, and E81-H226 are believed to stabilize the ATP bound conformation, while the decrease in the A176-I207 interaction may represent a shift in the NBD lobe positions towards a more ADP-bound conformation. This unbalanced induction of the ATP-bound conformation by JdH33Q may therefore serve as a basis for its inability to induce ATP hydrolysis in DnaK. Several other allosteric changes in intramolecular contacts were observed with JdH33Q binding. A significant increase in occupancy was seen for D148-Q442 (66.5% to 73.5%) which connect NBD lobe I and the SBDβ. D148 is known to sense signals from the substrate through residues V440 and L484, and trigger the release of SBDβ from the NBD (10). The observed change for with the association of D148 to the V440-proximal Q442 may have functional consequences. This increase in occupancy can support the prevention of NBD – SBD undocking and contribute to the inhibition of ATP hydrolysis.

### Interaction network between the Jd HPD and the DnaK catalytic center

Using the interaction data from the equilibration simulations, the shortest path from Jd to the catalytic center of the NBD was traced. Only interactions with occupancy greater than 30% were considered. The starting point was chosen as either D35/N35 and the end point was chosen as K70. In the wild-type and D35N mutant, the shortest path obtained was the same although the occupancies of each interaction within the path differ among samples. The path consists of D35/N35, R167, I168, I140, L9 and K70. However, in the H33Q mutant, the L9-K70 H-bond had an occupancy of 11.17%. The occupancies are shown in Table 1 and the bond interaction network linking D35 / N35 and K70 in the three binding pairs are shown in Figure 5.

**Figure 5.**
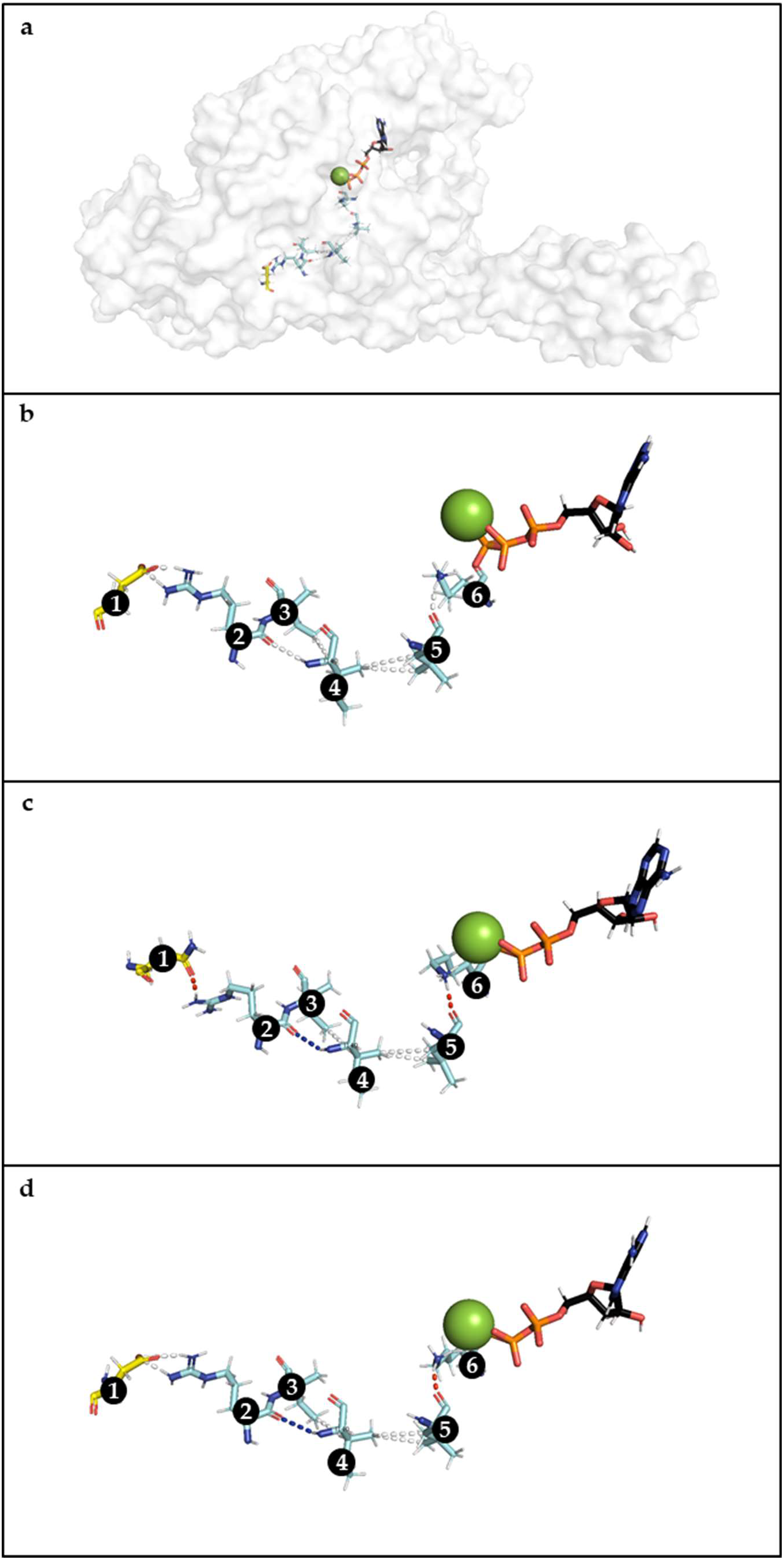
Path traced from Jd D35/N35 to DnaK K70. The shortest path from D35/N35 of Jd to the K70 at the catalytic center of DnaK was traced through the H-bonds, electrostatic and nonpolar interactions detected during equilibration. The figure shows the residues involved in the final frame of one replicate: (**a**) The position of the residues, ATP and Mg^2+^ within their respective molecules in the final frame of the simulation; (**b**) A close-up view of the residues in wildtype; (**c**) A close-up view of the residues in mutants D35N and (**d**) H33Q, colored based on changes in occupancy relative to wildtype (blue: decrease, red: increase, white: no change). Labels: (1) D35/N35, (2) R167, (3) I168, (4) I140, (5) L9, (6) K70

**Table 1.**
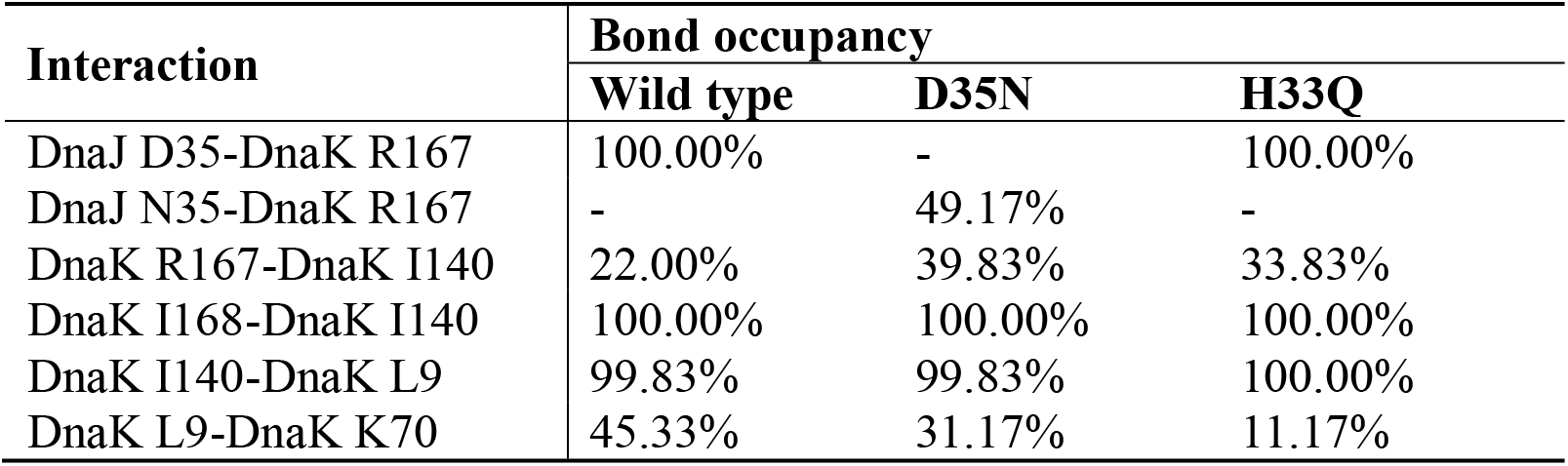
Residues involved in the path from Jd to the catalytic site.

| Interaction | Bond occupancy |  |  |
| --- | --- | --- | --- |
|  | Wild type | D35N | H33Q |
| DnaJ D35-DnaK R167 | 100.00% | - | 100.00% |
| DnaJ N35-DnaK R167 | - | 49.17% | - |
| DnaK R167-DnaK I140 | 22.00% | 39.83% | 33.83% |
| DnaK I168-DnaK I140 | 100.00% | 100.00% | 100.00% |
| DnaK I140-DnaK L9 | 99.83% | 99.83% | 100.00% |
| DnaK L9-DnaK K70 | 45.33% | 31.17% | 11.17% |

In the wildtype, D35 and R167 maintains a salt bridge throughout the whole simulation. R167 also forms an H-bond with I140 with an occupancy of 22%. However, its neighbor, I168 maintains a nonpolar interaction with I140, which could propagate the signal from Jd. I140 also forms a nonpolar interaction with L9 which finally reaches K70 through an H-bond with 45.33% occupancy.

In D35N, the conversion of D35 to N35 abolishes the salt bridge with R167. An H-bond of lower occupancy 49.17% is formed between N35 and R167. The H-bond between R167 and I140 in turn increased to 39.83% from 22% in wildtype. The nonpolar interactions between I140 and L9 remain at 99.83% but the H-bond occupancy between L9 and K70 decreases to 31.17%. The decreased occupancy for both the N35-R167, and the L9-K70 interactions suggest the decreased efficiency of JdD35N to promote ATP hydrolysis in the DnaK catalytic site.

In H33Q, the salt bridge between D35 and R167 was maintained as in wildtype. H-bonding between R167 and I140 was stronger at 33.83% occupancy and in addition, I168 maintains a nonpolar interaction with I140 and I140 in turn to L9. However, the H-bond occupancy between L9 and K70 in H33Q was decreased to 11.17%. Disruption of the direct link from D35 to K70, even at the last linkage (L9-K70), may be expected to cause a lower efficiency for promoting ATP hydrolysis in the DnaK catalytic site by JdH33Q. In addition, the decrease in occupancy for L9-K70 in both D35N and H33Q may cause the formation of a salt bridge between K70 and D8 in the mutants. This positions the stabilizing K70 farther from the active site, effectively decreasing ATPase activity.

We must note that residues I168 and I140 were not documented as part of the interaction network defined by Kityk et al. (9), however, these residues lie on a similar surface, and in adjacent beta sheets as the N170 residue indicated in the crystallized Hsp40-Hsp70 structure (9). Mutation of this residue has been observed to greatly affect the interaction of these two chaperone proteins (6). Our results suggest that modification of other residues proximal to N170 may also result in the modulation of Hsp40-Hsp70 interaction.

## CONCLUSION

The elucidation of a crystal structure for a whole DnaK-Jd complex (9) allows more detailed investigations of the system’s mechanism. In this study, molecular dynamics simulations were used to investigate the effects of the well-documented Jd dysfunctional mutants (D35N and H33Q) on DnaK function. By comparing the residue RMSDs of each mutant complex against the wildtype version, areas of increased mobility were identified around the active site of DnaK, and in the attached J-domains.

The mutations in the conserved Hsp40 HPD tripeptide resulted in changes for J-domain flexibility (Figure 1) and allosteric changes in the partner Hsp70 (Figures 2, 3 and 4). Binding of the mutant Jd forms was also observed to alter DnaK NBD lobe interactions (Figure 4).

The effect of the mutations was further evaluated based on observed occupancies of H-bonds, electrostatic and hydrophobic interactions. Changes in bond occupancy were observed in the contact surface between DnaK and Jd, between the two lobes of the DnaK NBD, and between the DnaK NBD and SBD. These observations suggest mechanisms through which the mutations on the conserved HPD tripeptide result in the induction of allosteric effects within DnaK that disrupt the promotion of ATP hydrolysis.

Changes in bond occupancy were also observed to involve the Hsp70 active site residues, including K70. The K70A mutation in DnaK has been previously observed to abolish ATPase activity and disrupt ATP-dependent conformational changes in DnaK (11). The shortest path from the HPD tripeptide to the active site was traced through 6 residues: D35/N35, R167, I168, I140, L9 and K70.

A network of important interactions between the Hsp40 Jd, the Hsp70 ATPase catalytic site, and the Hsp70 substrate recognition site has been previously constructed (9). The path traced in this study was similar to the previous report in 1 out of 4 interactions. Only D35-R167 was similar, while R167/I168-I140, I140-L9, and L9-K70 were not part of the previously proposed network. Our observed paths did not include the P143-K70, and A144-R15 links observed in their network.

The selected signaling path (6 residues) for this project was shorter by one residue than the shortest path available from the HPD tripeptide to the bound ATP in the previous report (7 residues).

The observed difference was based on the selection of the shortest path through hydrophobic interactions. For this study, a nonpolar interaction wassimply defined as interactions between hydrophobic residues (A, L, V, I, P, F, M, W) within 4 Å. The previous study considered several interaction modes connecting the D35 to the bound ATP, the shortest of which involved several bond types, and were less direct than the path observed with the current prodictions (9). This observed difference, as well as others, may be attributed to varied predictions of molecular movement and interactions in the simulated system due to the absence of the introduced disulfide linkage; unlike in the crystallized Jd-DnaK complex. This result supports the capacity of *in silico* methods to refine predictions on residue interaction networks, particularly for molecules whose structural elucidation *in vitro* requires specialized modifications.

The recent work by Tomiczek et al. (4) reveals altered interactions between Hsp40s and Hsp70s from different species. Despite the alterations, key residue contacts (i.e. R167-D393 and R167-D481) are kept consistent for these functional Hsp40-Hsp70 complexes. Our current data reveals changes in these key residue contacts for the dysfunctional mutant complexes. In particular, the interaction network linking the Jd HPD tripeptide with the DnaK active site was observed to deviate in the case of the Jd mutants. Key residues with altered bond occupancy included R167, D326, D148, and Q378. The JdD35N-induced increase in bond occupancy for R167 and D479 may represent a dysfunctional DnaK conformation induced by this mutant Jd form. Removal of this electrostatic interaction through an alanine substitution would be consistent with the observed rescue of the JdD35N defect by DnaK R167A (6).

The current work involved simulations of 0.1ns length. Interestingly, similar key residue contacts observed in these simulations, R167-D393, R167-D481, and R167-D35, were documented to persist in longer simulation runs (500ns) (4). These highlight crucial interactions whose impact on Hsp40-Hsp70 function are apparent even with short simulation times. These findings also suggest that divergence in functional mechanisms may be reliably predicted with relatively low computational cost, particularly for mutations with experimentally validated phenotypic significance. For this study, the observed divergence in the interaction network suggests a difference in the mechanisms involved between Hsp70 and the wildtype and mutant forms of Hsp40. Coupled with the altered interaction pair predominance, the data suggests that the dysfunction caused by the D35N and H33Q mutations propagates through changes in both intermolecular binding modes, as well as allosteric changes in intramolecular contacts that inhibit NBD and SBD undocking, and disrupt the efficient signal transfer from the conserved HPD tripeptide in Hsp40 to the catalytic residues in the ATPase domain, and the interdomain linker of Hsp70.

While the current manuscript focuses on the identification of key residues in the interaction between Hsp40 and Hsp70. This information may be used to guide future experiments, both *in silico* and *in vitro,* that would define an interaction network that may be targeted to control functions in other molecular chaperone machines. Moreover, the data reported illustrates how the applied methodology may be used to investigate the interaction networks between other protein binding partners and identify key residues for their modulation. This methodology may be utilized to target potential sites for correcting protein dysfunctions, and for optimizing protein functions for different industrial applications.

## AUTHOR CONTRIBUTIONS

*OOM: Investigation, Data Analysis, Manuscript Preparation and Revision*

*NADB: Conceptualization, Data Analysis, Manuscript Revision*

## ACKNOWLEDGEMENTS

The authors would like to thank Dr. Iris Ivy M. Gauran of the University of the Philippines School of Statistics for her advice regarding proper statistical testing. NADB would like to thank the University of the Philippines for continued support, allowing sustained work on this project despite the restrictions put for community lockdown.

